# BABAPPASnake: a workflow for episodic selection analysis with robustness-aware summaries

**DOI:** 10.1101/2025.04.27.650835

**Authors:** Samrat Singha, Palash Panda, Aloke Panda, Sanjib Kumar Das, Anumita Das, Nabanita Ghosh, Krishnendu Sinha

## Abstract

Episodic selection analyses are often assembled from fragmented toolchains in which ortholog discovery, codon alignment, phylogeny, exploratory scans, branch-site testing, and reporting are handled separately, making reproducibility and sensitivity tracking difficult. We introduce **BABAPPASnake** as an integrated workflow contribution for orthogroup-centered episodic selection analysis. The workflow combines orthogroup construction logic, CDS quality-aware mapping, multi-engine alignment pathways, phylogenetic inference, exploratory nomination, and branch-site follow-up testing in one reproducible execution framework. It also supports optional HyPhy GARD recombination screening as a conservative preprocessing report layer without forcing fragment-level rerouting by default. It generates pathway-level and cross-pathway robustness outputs, including matrix, consensus, narrative, and provenance summaries to support sensitivity-aware interpretation. A four-gene mosquito melanization-associated module is analyzed as a real-data empirical demonstration of end-to-end workflow behavior. In this demonstration, branch/site signals show both recurrent and method-sensitive components across six method-trim pathways, with a directional core-tier tendency in several summaries. These case-study patterns are interpreted as workflow-based empirical evidence and hypothesis-generating asymmetry, not decisive pathway-level confirmation. Overall, BABAPPASnake provides a practical and reproducible framework for episodic selection studies where analytical sensitivity must be explicitly reported.

## 1 Introduction

Episodic selection analysis is widely used in molecular evolution, but practical implementation remains technically fragmented. Users typically assemble separate tools for ortholog collection, alignment, trimming, tree reconstruction, exploratory scans, branch-site testing, and post hoc reporting. This fragmentation creates reproducibility risks because important analytical decisions are distributed across scripts, formats, and manual steps.

A second challenge is analytical sensitivity. Branch-level and site-level outcomes can shift across orthogroup selection rules, alignment engines, trim states, and downstream model settings. When this sensitivity is not reported explicitly, conclusions can be overinterpreted as if they were workflow-invariant.

BABAPPASnake was developed to address these practical issues by integrating major episodic-selection stages into one workflow with organized outputs and explicit robustness summaries. The goal is not to claim replacement of established inference engines, but to combine them in a reproducible structure that exposes both stable and method-sensitive signals.

As an empirical demonstration context, we use a four-gene mosquito melanization-associated module with biologically relevant core and catalytic pathway tiers [Kanost and Jiang, 2015, Povelones et al., 2013, El Moussawi et al., 2019, Sousa et al., 2020, Saab et al., 2024, Morejon and Michel, 2025]. In this manuscript, biology is presented as a real-data use case for workflow utility, not as the sole endpoint claim.

### Contribution statement

This study introduces BABAPPASnake as a robustness-aware episodic selection workflow and demonstrates its practical use on a mosquito melanization-associated gene module.

## 2 BABAPPASnake Workflow Design

### 2.1 Architecture and rule order

BABAPPASnake organizes analysis around a query-centered, rule-ordered workflow rather than a single black-box call. The operational logic in the current implementation is:

1. read query/proteome inputs and run orthogroup discovery through RBH and OrthoFinder routes;
2. map the query to OrthoFinder orthogroups by BLASTP against orthogroup-member proteins (not query-ID membership), then compare strict 1:1 ortholog counts and choose the better backend (RBH on ties);
3. if both backends produce zero strict 1:1 support, stop explicitly; if CDS is not yet supplied, emit `WAITING_FOR_CDS.txt` and stop for staged resume;
4. if CDS is supplied, run CDS mapping and quality control (lowercase intron clipping, ORF/frame checks, and mapped CDS/protein output);
5. expand to method*×*trim pathways (selected methods: BABAPPAlign, MAFFT, PRANK; forced trim states: raw+ClipKIT);
6. build protein/codon alignments, run IQ-TREE per pathway, apply optional outgroup rooting, optionally run HyPhy GARD recombination screening, run HyPhy exploratory scans, parse dynamic foregrounds, run branch-site codeml follow-up, run codeml ASR, and then extract selected-branch ancestral/descendant sequence changes;
7. write pathway summaries, cross-pathway robustness outputs, top-level aliases, and machine-readable run provenance.

Core inputs are a query protein FASTA, a proteome directory, and an optional CDS FASTA/outgroup string. The workflow supports guided interactive execution and non-guided execution, with resumable stage-wise behavior.

### 2.2 Orthogroup, CDS, and alignment pathways

In the default route, RBH and OrthoFinder outputs are compared, and downstream analysis uses the better strict 1:1 ortholog support set. CDS records are reconciled to orthogroup proteins with quality filters (including intron-clipping and ORF checks). Alignment pathways include BABAPPAlign [Sinha, 2025], MAFFT [Katoh and Standley, 2013], and PRANK [Löytynoja, 2021]; each is evaluated under raw and ClipKIT trim states. Thus, one run can generate six pathway branches (method*×*trim).

### 2.3 Inference and follow-up logic

Per pathway, BABAPPASnake executes IQ-TREE phylogeny estimation [Wong et al., 2025, Kalyaanamoorthy et al., 2017], optional outgroup rooting, HyPhy aBSREL/MEME exploratory scans [Kosakovsky Pond et al., 2020], and branch-site codeml follow-up testing [Yang, 2007, Álvarez-Carretero et al., 2023]. Foreground nomination is exploratory, while branch-site codeml is treated as follow-up testing with within-gene BH correction.

### 2.4 Optional recombination-screening branch

BABAPPASnake now includes an optional HyPhy GARD recombination-screening module that can be enabled per run. When enabled, GARD is executed per active method*×*trim pathway and writes pathway-specific raw JSON, logs, and parsed summary metadata. In the current implementation, this branch is conservative: it reports breakpoint evidence and pathway-level recombination status but does not automatically reroute downstream branchsite analyses onto recombinant fragments. Accordingly, full-length pathway inference remains the default unless explicit fragment-aware routing is added in a future release.

### 2.5 Robustness and reproducibility outputs

The workflow writes pathway-specific and cross-pathway outputs, including branch reproducibility matrices, pathway-tier summaries, and narrative/consensus robustness reports. Machine-readable provenance and structured output directories support archive-oriented reproducibility and review.

### 2.6 Software use: installation and execution

#### Dependencies installed separately (external executables)

`blastp, makeblastdb, orthofinder, iqtree` (or `iqtree2/iqtree3), hyphy, codeml` (PAML), `clipkit`; optional but recommended: `mafft, prank`.

#### Python package installation

~~~
conda create -n babappasnake -c conda-forge -c bioconda \
 python=3.11 blast orthofinder iqtree hyphy paml clipkit mafft prank pip
conda activate babappasnake
pip install babappasnake
~~~

#### Example run (non-interactive)

~~~
babappasnake \
 --prot /path/to/proteomes \
 --query /path/to/query.fasta \
 --cds /path/to/orthogroup_cds.fasta \
 --alignment-methods 4 \
 --outgroup culex \
 --outdir run01 \
 --threads 12 \
 --interactive no \
 --guided no
~~~

#### Example run (guided interactive mode)

~~~
babappasnake
~~~

Interactive mode executes one stage at a time with run/skip/stop control and safe resume on rerun.

## 3 Results

### 3.1 Workflow outputs and analytical organization

The workflow produced organized outputs across six pathways (babappalign, MAFFT, and PRANK; each under raw and ClipKIT) for each gene. Branch-level comparability was retained across all pathways, while site-level comparability was partial in pathways with recorded MEME failures for CLIPA8. This output design made it possible to separate recurrent branch-level patterns from method-sensitive outcomes. The current software release additionally supports optional per-pathway GARD recombination screening summaries; these are reported as preprocessing evidence and do not alter the default full-length branch-site execution chain.

**Figure 1:**
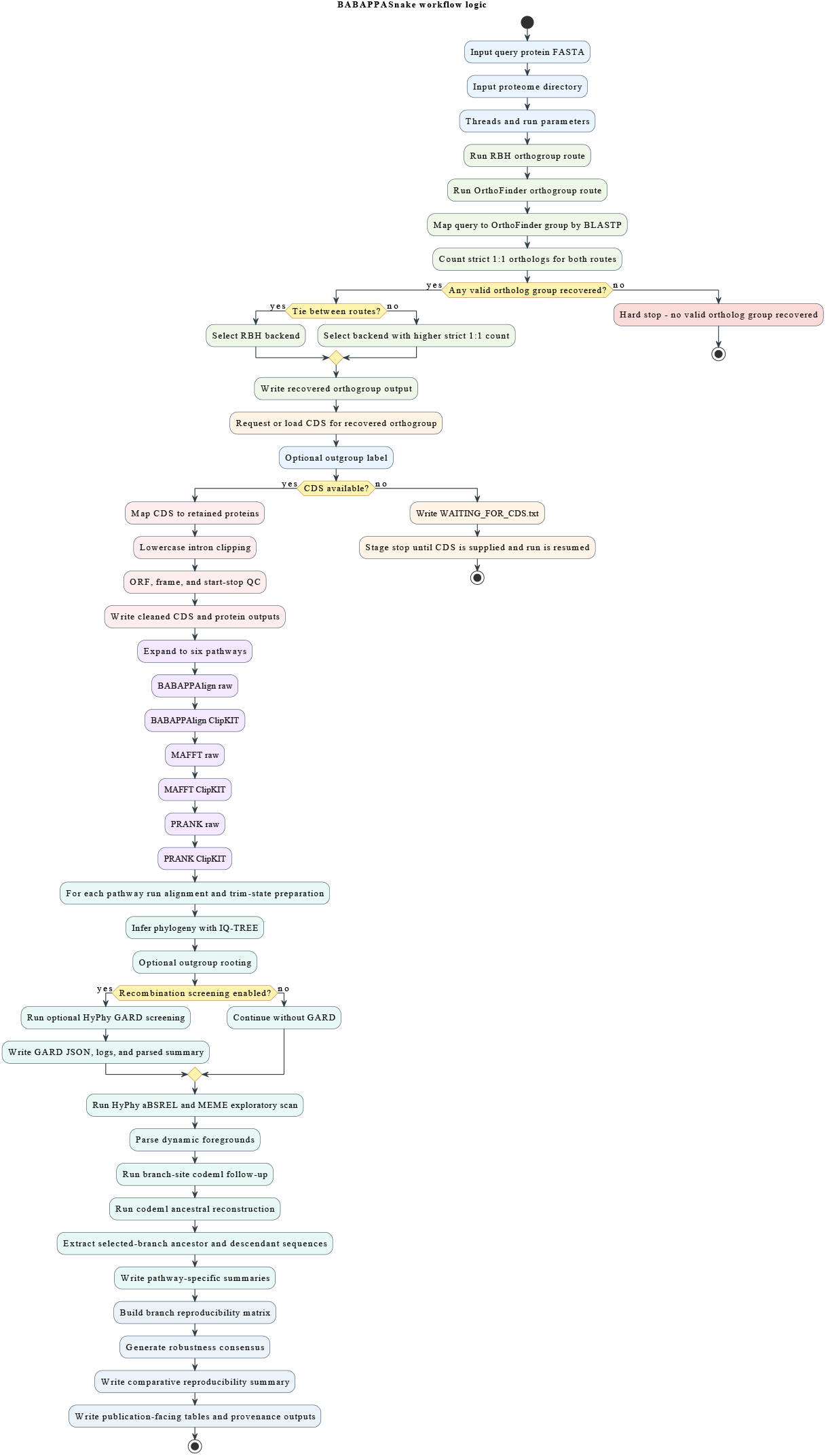
Decision-aware BABAPPASnake architecture. The workflow starts with RBH and OrthoFinder orthogroup discovery, applies strict 1:1 backend selection (tie *→* RBH), and stops explicitly when both routes return zero support. A CDS stage gate enables staged stop/resume execution before expanding into six method *×*trim pathways that each follow the same inference chain (alignment, tree inference/rooting, optional GARD screening, HyPhy exploration, branch-site follow-up, and ASR extraction). Pathway outputs then converge into cross-pathway robustness summaries and run-level provenance outputs for reproducible interpretation.

Table 1 summarizes key workflow-level behavior from the demonstration, including pathway totals and reproducibility classes.

**Table 1:**
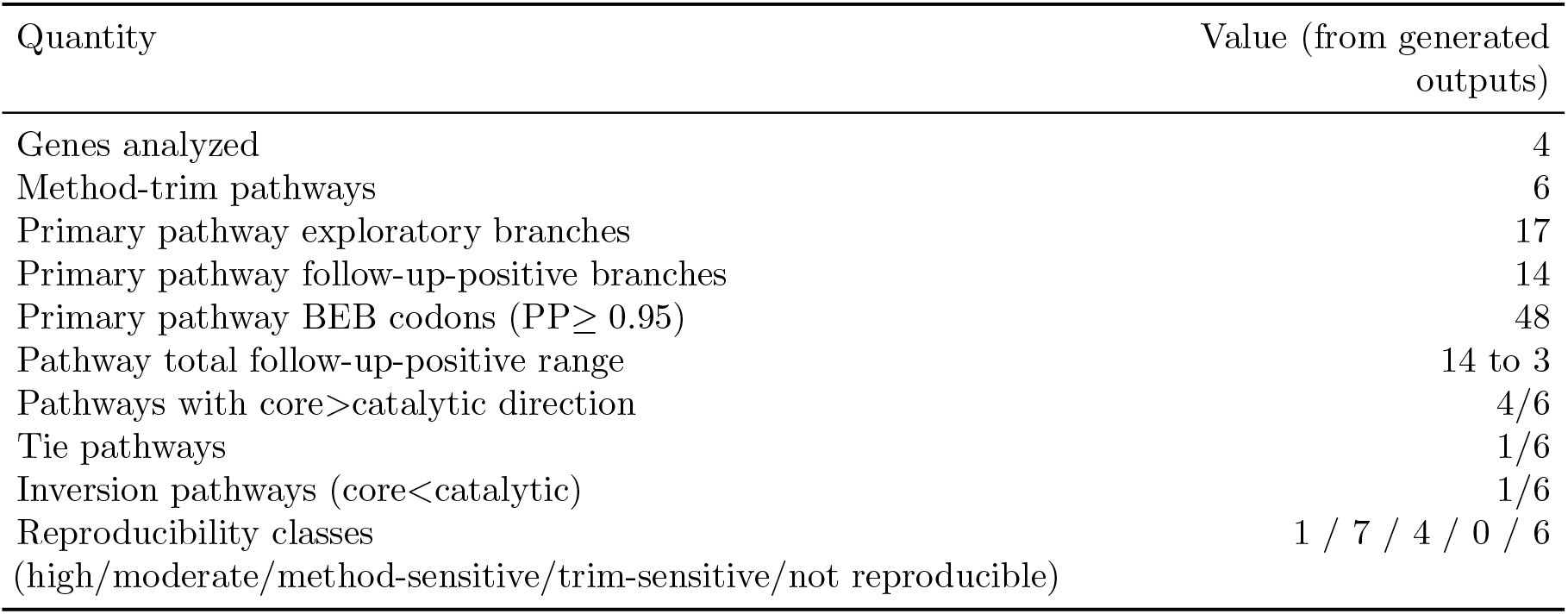
Workflow-level summary of empirical demonstration outputs.

### 3.2 Empirical mosquito case study as workflow demonstration

In the predefined primary pathway (babappalign+ClipKIT), 17 exploratory aBSREL candidates yielded 14 branch-site follow-up-positive branches after BH correction. The complement-coupled core tier (*SPCLIP1* +*CLIPA8*) contributed 11/14 follow-up-positive branches, while the catalytic tier (*CLIPB14* +*CLIPB15*) contributed 3/14. At site level, 48 BEB codons (PP*≥* 0.95) were identified, with 43 in the core tier and 5 in the catalytic tier.

Global one-ratio codeml values remained in purifying-selection range (*ω* = 0.186 to 0.297), supporting interpretation as episodic signals on constrained backgrounds. Across six pathways, the directional split recurred in 4/6 pathways (one tie, one inversion), indicating a suggestive asymmetry pattern but non-uniform branch-level stability.

Exact asymmetry tests remained non-significant in the current dataset (one-sided *p* = 0.191, 0.167, and 0.188). Accordingly, the demonstration is interpreted as directional and hypothesis-generating rather than statistically decisive.

## 4 Discussion

The main contribution of this paper is methodological: BABAPPASnake provides an integrated, reproducibility-oriented workflow for episodic selection analyses that are usually fragmented in practice. By presenting method*×*trim pathway outputs and explicit cross-pathway summaries, it enables robustness-aware interpretation rather than single-path overstatement.

The mosquito four-gene analysis serves as an empirical end-to-end demonstration of utility. It shows that the workflow can recover recurring patterns while simultaneously exposing sensitivity and fragility where they exist. This dual reporting is a practical advantage for downstream biological prioritization.

The case-study biology is therefore interpreted as a real-data illustration of workflow behavior. Observed pathway-tier asymmetry is treated as a suggestive directional trend and hypothesis-generating pattern, not as decisive enrichment.

### Limitations and future validation

This manuscript does not provide broad bench-marking against alternative end-to-end pipelines. That remains future work. Additional future directions include expanded orthology stress-testing, fragment-aware downstream routing after GARD screening, and structural/domain mapping of candidate codons.

## Code availability

BABAPPASnake repository: https://github.com/sinhakrishnendu/babappasnake.git The repository includes an optional HyPhy GARD recombination-screening module (‘–recombination’) with pathway-level reporting outputs.

## Data/archive availability

Case-study run outputs, generated tables, and figure assets used for this manuscript are available in the archived study bundle (https://doi.org/10.5281/zenodo.19048331).

## Supplementary information

Supplementary material provides workflow details, case-study method specifics, robustness/reproducibility tables, supplementary figures, and archive file-guide mapping.

## Reproducibility statement

The software repository provides reusable workflow implementation; the archived run bundle provides case-study-specific outputs and generated manuscript tables/figures; the supplementary file provides curated manuscript-facing organization of those outputs.

## Acknowledgements

NG and KS are deeply grateful to their little son Shaswata, whose boundless curiosity and love for butterflies inspired the name “BABAPPA.” In his imaginative vocabulary, “BABAPPA” refers to a butterfly, and it was this creativity that ultimately gave the software its name. We thank the open-source bioinformatics community for tools and resources used in this workflow.

