## Supplementary material for "BABAPPASnake: a workflow for episodic selection analysis with robustness-aware summaries"

### Supplementary Material
### BABAPPASnake Workflow Paper Reform

#### Supplementary Workflow Details

##### S1. Workflow scope

BABAPPASnake is used here as an integrated workflow for orthogroup-centered episodic selection analysis. The manuscript-level implementation includes orthogroup definition, CDS mapping/QC, multi-method codon-aware alignment pathways, IQ-TREE phylogeny, HyPhy exploratory nomination, branch-site codeml follow-up testing, and robustness-oriented summaries.

##### S2. Input and pathway structure

The empirical demonstration uses four genes in a mosquito melanization-associated module. For each gene, six pathways were run:

- babappalign_raw
- babappalign_clipkit
- mafft_raw
- mafft_clipkit
- prank_raw
- prank_clipkit

This method×trim branching is central to the workflow's robustness-aware design.

##### S3. Follow-up testing hierarchy

The inferential hierarchy used throughout is:

1. exploratory branch nomination by aBSREL;
2. branch-site codeml follow-up testing for nominated branches;
3. within-gene BH correction for follow-up calls.

In this study, "follow-up" does not imply decisive confirmation; results are interpreted with explicit pathway sensitivity context.

#### S4. Optional recombination-screening module

BABAPPASnake includes an optional HyPhy GARD screening branch that can be enabled per run. In the current implementation, this module is conservative: it records pathway-level recombination evidence and breakpoint reporting metadata, but downstream branch-site analyses remain full-length by default. This preserves backward compatibility while exposing recombination evidence explicitly in reproducibility summaries.

### Supplementary Case Study Methods

#### S5. Primary summary values used in manuscript

All values below are taken from generated tables already included with the manuscript project:

- Primary pathway exploratory candidates: 17
- Primary pathway follow-up-positive branches: 14
- Core-tier follow-up-positive branches: 11
- Catalytic-tier follow-up-positive branches: 3
- Total BEB codons (PP$\geq$ 0.95): 48
- Core-tier BEB codons: 43
- Catalytic-tier BEB codons: 5
- One-ratio $\omega$ range: 0.186–0.297
- Pathway direction summary: core>catalytic in 4/6, one tie, one inversion
- Exact asymmetry tests (one-sided): $p = 0.191$, 0.167, 0.188

#### S6. Statistical framing used in revised manuscript

The revised manuscript treats these values as a directional and hypothesis-generating pattern in a workflow demonstration. The asymmetry is explicitly reported as non-significant at conventional thresholds, and biological claims are presented cautiously.

Table 1: Primary pathway summary by gene (babappalign+ClipKIT).

| Gene | Tier | $\omega$ | aBSREL candidates | Follow-up positive | BEB codons | MEME sites |
|---|---|---|---|---|---|---|
| CLIPA8 | core | 0.24527 | 6 | 6 | 23 | 10 |
| CLIPB14 | catalytic | 0.24464 | 3 | 2 | 3 | 8 |
| CLIPB15 | catalytic | 0.18612 | 2 | 1 | 2 | 6 |
| SPCLIP1 | core | 0.29731 | 6 | 5 | 20 | 12 |

Table 2: Core vs catalytic follow-up-positive branch counts by pathway.

| Pathway | Core confirmed | Catalytic confirmed | Direction |
|---|---|---|---|
| babappalign_raw | 8 | 3 | core>catalytic |
| babappalign_clipkit | 11 | 3 | core>catalytic |
| mafft_raw | 5 | 3 | core>catalytic |
| mafft_clipkit | 2 | 2 | tie |
| prank_raw | 3 | 2 | core>catalytic |
| prank_clipkit | 1 | 2 | catalytic>core |

Table 3: Lineage reproducibility classes across six pathways.

| Class | Number of lineages |
|---|---|
| highly_robust | 1 |
| moderately_robust | 7 |
| method_sensitive | 4 |
| trim_sensitive | 0 |
| not_reproducible | 6 |

### Supplementary Tables

#### S7. Primary gene-level summary

#### S8. Six-pathway tier summary

#### S9. Reproducibility class totals

### Robustness and Reproducibility Outputs

#### S10. Full content of generated TSV files

Each generated TSV used for interpretation is reproduced below in full.

##### S10.1 primary_gene_summary.tsv

```
gene query_id tier rbh_orthogroup_members cds_mappings_retained final_alignment_sequences
    final_codon_sites omega_one_ratio absrel_candidates codeml_confirmed beb_sites_branch_summed_pp95
    meme_sites_p_le_0.05 meme_beb_overlap_count meme_beb_overlap_sites
CLIPA8 AGAP010731 core 31 29 29 331 0.24527 6 6 23 10 2 11,67
CLIPB14 AGAP010833 catalytic 31 30 30 308 0.24464 3 2 3 8 0
CLIPB15 AGAP009844 catalytic 31 30 30 307 0.18612 2 1 2 6 2 146,216
SPCLIP1 AGAP028725 core 30 30 30 347 0.29731 6 5 20 12 1 336
```

##### S10.2 confirmed_branchsite_table.tsv

```
gene query_id tier foreground_branch canonical_lineage display_lineage absrel_p lrt codeml_p codeml_q
    confirmed_bh_0.05 beb_site_count_pp95 beb_sites_pp95
CLIPA8 AGAP010731 core AFUN2_002201_R2179_Anopheles_funestus_AfunGA1 funestus A. funestus 0.0
    57.98460999999952 2.6417649450312327e-14 1.5850589670187396e-13 True 5 4W, 11V, 47Y, 67M, 173R
CLIPA8 AGAP010731 core AMOUCM1_004393_R4353_Anopheles_moucheti_AmouCM1 moucheti A. moucheti
    0.0003840568736150907 9.651993999999831 0.0018914615542106293 0.002269753865052755 True 3 72E, 166V,
    239Y
CLIPA8 AGAP010731 core ACRUBR1_007112_R7575_Anopheles_cruzii_AcruBR1 cruzii A. cruzii 0.00467980614794028
    23.483507999997528 1.259891359301696e-06 3.7796740779050885e-06 True 7 49P, 70F, 72E, 189P, 223D, 273
    E, 279D
CLIPA8 AGAP010731 core AMAC_004066_R3862_Anopheles_maculipalpis_AmacGA1 maculipalpis A. maculipalpis
    0.01705260478237969 5.114082000000053 0.023732419096545 0.023732419096545 True 0 None
CLIPA8 AGAP010731 core ACHR004918_RA_Anopheles_christyi_ACHKN1017 christyi A. christyi 0.02005301994149333
     13.632957999994687 0.00022224945079261704 0.0004444989015852341 True 6 50N, 65I, 77Y, 162T, 250R,
    321T
CLIPA8 AGAP010731 core ADAR2_010765_R17226_Anopheles_darlingi_AdarGF1 darlingi A. darlingi
    0.04094777785808468 10.233850000004168 0.001378865536846588 0.002068298305269882 True 2 122N, 153P
CLIPB14 AGAP010833 catalytic AMAC_000985_R425_Anopheles_maculipalpis_AmacGA1 maculipalpis A. maculipalpis
    2.528387648248476e-05 7.795996000000741 0.005236213853260576 0.007854320779890865 True 0 None
CLIPB14 AGAP010833 catalytic AMIN006979_RA_Anopheles_minimus_MINIMUS1 minimus A. minimus
    0.002021896244924049 23.373759999998583 1.3338568938849646e-06 4.001570681654894e-06 True 3 150M, 246
    K, 308A
CLIPB15 AGAP009844 catalytic AMOUCM1_000648_R17196_Anopheles_moucheti_AmouCM1 moucheti A. moucheti
    0.002178620267505038 12.911605999997846 0.00032655098621746647 0.0006531019724349329 True 2 146S, 216
    R
SPCLIP1 AGAP028725 core ACUA011631_RA_Anopheles_culicifacies culicifacies A. culicifacies
    4.500256667316904e-08 34.571888000005856 4.107962644759239e-09 2.4647775868555437e-08 True 6 106T,
    113P, 114D, 208A, 225F, 336R
SPCLIP1 AGAP028725 core AFUN2_012038_R21717_Anopheles_funestus_AfunGA1 funestus A. funestus
    0.00100068482323673 11.431797999997798 0.0007219778667420193 0.0011983233246788435 True 5 206V, 238N,
     240P, 252A, 273L
```

```
SPCLIP1 AGAP028725 core AMEC000258_RA_Anopheles_melas_CM1001059_A melas A. melas 0.002325502954313885 9.016788000000815 0.002675110727269404 0.003210132872723285 True 0 None
SPCLIP1 AGAP028725 core ACRUBR1_005064_R5104_Anopheles_cruzii_AcruBR1 cruzii A. cruzii 0.006199146895861818 11.243827999998757 0.0007988822164525624 0.0011983233246788435 True 1 336R
SPCLIP1 AGAP028725 core AMIN004295_RA_Anopheles_minimus_MINIMUS1 minimus A. minimus 0.007316637533225978 22.94249200000195 1.6692088405716758e-06 5.007626521715028e-06 True 8 164R, 212L, 238N, 240P, 243N, 265T, 281Y, 307A
```

#### S10.3 exploratory_not_confirmed.tsv

```
gene query_id foreground_branch canonical_lineage display_lineage absrel_p codeml_p codeml_q
CLIPB14 AGAP010833 ADAR2_004445_R6063_Anopheles_darlingi_AdarGF1 darlingi A. darlingi 0.00196688085224328 1.0 1.0
CLIPB15 AGAP009844 AMIN009569_RA_Anopheles_minimus_MINIMUS1 minimus A. minimus 0.003398615876087585 0.19022217812820444 0.19022217812820444
SPCLIP1 AGAP028725 AMAM010672_RA_Anopheles_maculatus_maculatus3 maculatus A. maculatus 0.04490798880698171 1.0 1.0
```

#### S10.4 alignment_robustness_matrix.tsv

```
gene query_id tier pathway method trim_state status absrel_status meme_status branch_level_interpretable site_level_interpretable absrel_threshold meme_threshold n_sequences protein_alignment_length codon_alignment_length_nt n_absrel_significant n_branchsite_significant_after_bh
CLIPA8 AGAP010731 core babappalign_raw babappalign raw hyphy_failed ok failed True False 0.05 0.05 29 532 1491 5 4
CLIPA8 AGAP010731 core babappalign_clipkit babappalign clipkit ok ok ok True True 0.05 0.05 29 358 993 6 6
CLIPA8 AGAP010731 core mafft_raw mafft raw hyphy_failed ok failed True False 0.05 0.05 29 600 1800 4 3
CLIPA8 AGAP010731 core mafft_clipkit mafft clipkit ok ok ok True True 0.05 0.05 29 356 843 1 1
CLIPA8 AGAP010731 core prank_raw prank raw ok ok ok True True 0.05 0.05 29 924 2772 2 2
CLIPA8 AGAP010731 core prank_clipkit prank clipkit hyphy_failed ok failed True False 0.05 0.05 29 360 945 1 1
CLIPB14 AGAP010833 catalytic babappalign_raw babappalign raw ok ok ok True True 0.05 0.05 30 418 1239 4 2
CLIPB14 AGAP010833 catalytic babappalign_clipkit babappalign clipkit ok ok ok True True 0.05 0.05 30 349 924 3 2
CLIPB14 AGAP010833 catalytic mafft_raw mafft raw ok ok ok True True 0.13 0.05 30 444 1332 1 1
CLIPB14 AGAP010833 catalytic mafft_clipkit mafft clipkit ok ok ok True True 0.2 0.05 30 351 927 0 0
CLIPB14 AGAP010833 catalytic prank_raw prank raw ok ok ok True True 0.09 0.05 30 502 1506 1 1
CLIPB14 AGAP010833 catalytic prank_clipkit prank clipkit ok ok ok True True 0.2 0.05 30 371 942 1 1
CLIPB15 AGAP009844 catalytic babappalign_raw babappalign raw ok ok ok True True 0.05 0.05 30 409 1227 2 1
CLIPB15 AGAP009844 catalytic babappalign_clipkit babappalign clipkit ok ok ok True True 0.05 0.05 30 348 921 2 1
CLIPB15 AGAP009844 catalytic mafft_raw mafft raw ok ok ok True True 0.05 0.05 30 409 1227 2 2
CLIPB15 AGAP009844 catalytic mafft_clipkit mafft clipkit ok ok ok True True 0.05 0.05 30 352 693 2 2
CLIPB15 AGAP009844 catalytic prank_raw prank raw ok ok ok True True 0.05 0.05 30 465 1395 1 1
CLIPB15 AGAP009844 catalytic prank_clipkit prank clipkit ok ok ok True True 0.05 0.05 30 365 714 1 1
SPCLIP1 AGAP028725 core babappalign_raw babappalign raw ok ok ok True True 0.05 0.05 30 515 1509 5 4
SPCLIP1 AGAP028725 core babappalign_clipkit babappalign clipkit ok ok ok True True 0.05 0.05 30 395 1041 6 5
SPCLIP1 AGAP028725 core mafft_raw mafft raw ok ok ok True True 0.05 0.05 30 531 1593 2 2
SPCLIP1 AGAP028725 core mafft_clipkit mafft clipkit ok ok ok True True 0.05 0.05 30 389 1041 2 1
SPCLIP1 AGAP028725 core prank_raw prank raw ok ok ok True True 0.09 0.05 30 660 1980 1 1
SPCLIP1 AGAP028725 core prank_clipkit prank clipkit ok ok ok True True 0.2 0.05 30 419 1056 0 0
```

#### S10.5 pathway_tier_summary.tsv

```
pathway core_absrel_candidates catalytic_absrel_candidates core_codeml_confirmed catalytic_codeml_confirmed core_exceeds_catalytic_confirmed status
babappalign_raw 10 6 8 3 True partial_meme_failure
babappalign_clipkit 12 5 11 3 True ok
```

```
mafft_raw 6 3 5 3 True partial_meme_failure
mafft_clipkit 3 2 2 2 False ok
prank_raw 3 2 3 2 True ok
prank_clipkit 1 2 1 2 False partial_meme_failure
```

##### S10.6 branch_reproducibility_matrix.tsv

```
gene tier canonical_lineage display_lineage raw_branch_labels replication_count reproducibility_class
    babappalign_raw babappalign_clipkit mafft_raw mafft_clipkit prank_raw prank_clipkit
CLIPA8 core christyi A. christyi ACHR004918_RA_Anopheles_christyi_ACHKN1017 2 method_sensitive 1 1 0 0 0 0
CLIPA8 core cruzii A. cruzii ACRUBR1_007112_R7575_Anopheles_cruzii_AcruBR1 3 moderately_robust 1 1 1 0 0 0
CLIPA8 core darlingi A. darlingi ADAR2_010765_R17226_Anopheles_darlingi_AdarGF1 1 not_reproducible 0 1 0 0
     0 0
CLIPA8 core funestus A. funestus AFUN021726_RA_Anopheles_funestus_FUMOZ;
    AFUN2_002201_R2179_Anopheles_funestus_AfunGA1 4 moderately_robust 1 1 1 0 1 0
CLIPA8 core maculipalpis A. maculipalpis AMAC_004066_R3862_Anopheles_maculipalpis_AmacGA1 4
    moderately_robust 1 1 1 1 0 0
CLIPA8 core moucheti A. moucheti AMOUCM1_004393_R4353_Anopheles_moucheti_AmouCM1 3 moderately_robust 0 1 0
     0 1 1
CLIPB14 catalytic atroparvus A. atroparvus AATE006733_RA_Anopheles_atroparvus_EBRO 1 not_reproducible 1 0
    0 0 0 0
CLIPB14 catalytic gambiae A. gambiae AGAP010833 1 not_reproducible 0 0 0 0 1 0
CLIPB14 catalytic maculipalpis A. maculipalpis AMAC_000985_R425_Anopheles_maculipalpis_AmacGA1 1
    not_reproducible 0 1 0 0 0 0
CLIPB14 catalytic minimus A. minimus AMIN006979_RA_Anopheles_minimus_MINIMUS1 4 moderately_robust 1 1 1 0
    0 1
CLIPB15 catalytic aquasalis A. aquasalis AAQUA_001018_R21657_Anopheles_aquasalis 2 method_sensitive 0 0 1
    1 0 0
CLIPB15 catalytic moucheti A. moucheti AMOUCM1_000648_R17196_Anopheles_moucheti_AmouCM1 6 highly_robust 1
    1 1 1 1 1
SPCLIP1 core aquasalis A. aquasalis AAQUA_009556_R14763_Anopheles_aquasalis 1 not_reproducible 0 0 1 0 0 0
SPCLIP1 core cruzii A. cruzii ACRUBR1_005064_R5104_Anopheles_cruzii_AcruBR1 2 method_sensitive 1 1 0 0 0 0
SPCLIP1 core culicifacies A. culicifacies ACUA011631_RA_Anopheles_culicifacies 3 moderately_robust 1 1 0 0
     1 0
SPCLIP1 core funestus A. funestus AFUN2_012038_R21717_Anopheles_funestus_AfunGA1 2 method_sensitive 1 1 0
    0 0 0
SPCLIP1 core melas A. melas AMEC000258_RA_Anopheles_melas_CM1001059_A 1 not_reproducible 0 1 0 0 0 0
SPCLIP1 core minimus A. minimus AMIN004295_RA_Anopheles_minimus_MINIMUS1 4 moderately_robust 1 1 1 1 0 0
```

##### S10.7 robustness_consensus_branchsite.tsv

```
gene query_id tier canonical_lineage display_lineage raw_branch_labels replication_count
    replication_fraction methods_present trim_states_present pathways_present n_partial_pathways
    partial_pathways reproducibility_class
CLIPA8 AGAP010731 core christyi A. christyi ACHR004918_RA_Anopheles_christyi_ACHKN1017 2 0.333 babappalign
     clipkit,raw babappalign_clipkit,babappalign_raw 1 babappalign_raw method_sensitive
CLIPA8 AGAP010731 core cruzii A. cruzii ACRUBR1_007112_R7575_Anopheles_cruzii_AcruBR1 3 0.500 babappalign,
    mafft clipkit,raw babappalign_clipkit,babappalign_raw,mafft_raw 2 babappalign_raw,mafft_raw
    moderately_robust
CLIPA8 AGAP010731 core darlingi A. darlingi ADAR2_010765_R17226_Anopheles_darlingi_AdarGF1 1 0.167
    babappalign clipkit babappalign_clipkit 0  not_reproducible
CLIPA8 AGAP010731 core funestus A. funestus AFUN021726_RA_Anopheles_funestus_FUMOZ;
    AFUN2_002201_R2179_Anopheles_funestus_AfunGA1 4 0.667 babappalign,mafft,prank clipkit,raw
    babappalign_clipkit,babappalign_raw,mafft_raw,prank_raw 2 babappalign_raw,mafft_raw moderately_robust
CLIPA8 AGAP010731 core maculipalpis A. maculipalpis AMAC_004066_R3862_Anopheles_maculipalpis_AmacGA1 4
    0.667 babappalign,mafft clipkit,raw babappalign_clipkit,babappalign_raw,mafft_clipkit,mafft_raw 2
    babappalign_raw,mafft_raw moderately_robust
CLIPA8 AGAP010731 core moucheti A. moucheti AMOUCM1_004393_R4353_Anopheles_moucheti_AmouCM1 3 0.500
    babappalign,prank clipkit,raw babappalign_clipkit,prank_clipkit,prank_raw 1 prank_clipkit
    moderately_robust
CLIPB14 AGAP010833 catalytic atroparvus A. atroparvus AATE006733_RA_Anopheles_atroparvus_EBRO 1 0.167
    babappalign raw babappalign_raw 0  not_reproducible
```

```
CLIPB14 AGAP010833 catalytic gambiae A. gambiae AGAP010833 1 0.167 prank raw prank_raw 0  not_reproducible
CLIPB14 AGAP010833 catalytic maculipalpis A. maculipalpis AMAC_000985_R425_Anopheles_maculipalpis_AmacGA1
    1 0.167 babappalign clipkit babappalign_clipkit 0  not_reproducible
CLIPB14 AGAP010833 catalytic minimus A. minimus AMIN006979_RA_Anopheles_minimus_MINIMUS1 4 0.667
    babappalign,mafft,prank clipkit,raw babappalign_clipkit,babappalign_raw,mafft_raw,prank_clipkit 0
    moderately_robust
CLIPB15 AGAP009844 catalytic aquasalis A. aquasalis AAQUA_001018_R21657_Anopheles_aquasalis 2 0.333 mafft
    clipkit,raw mafft_clipkit,mafft_raw 0  method_sensitive
CLIPB15 AGAP009844 catalytic moucheti A. moucheti AMOUCM1_000648_R17196_Anopheles_moucheti_AmouCM1 6 1.000
     babappalign,mafft,prank clipkit,raw babappalign_clipkit,babappalign_raw,mafft_clipkit,mafft_raw,
    prank_clipkit,prank_raw 0  highly_robust
SPCLIP1 AGAP028725 core aquasalis A. aquasalis AAQUA_009556_R14763_Anopheles_aquasalis 1 0.167 mafft raw
    mafft_raw 0  not_reproducible
SPCLIP1 AGAP028725 core cruzii A. cruzii ACRUBR1_005064_R5104_Anopheles_cruzii_AcruBR1 2 0.333 babappalign
     clipkit,raw babappalign_clipkit,babappalign_raw 0  method_sensitive
SPCLIP1 AGAP028725 core culicifacies A. culicifacies ACUA011631_RA_Anopheles_culicifacies 3 0.500
    babappalign,prank clipkit,raw babappalign_clipkit,babappalign_raw,prank_raw 0  moderately_robust
SPCLIP1 AGAP028725 core funestus A. funestus AFUN2_012038_R21717_Anopheles_funestus_AfunGA1 2 0.333
    babappalign clipkit,raw babappalign_clipkit,babappalign_raw 0  method_sensitive
SPCLIP1 AGAP028725 core melas A. melas AMEC000258_RA_Anopheles_melas_CM1001059_A 1 0.167 babappalign
    clipkit babappalign_clipkit 0  not_reproducible
SPCLIP1 AGAP028725 core minimus A. minimus AMIN004295_RA_Anopheles_minimus_MINIMUS1 4 0.667 babappalign,
    mafft clipkit,raw babappalign_clipkit,babappalign_raw,mafft_clipkit,mafft_raw 0  moderately_robust
```

#### S10.8 reproducibility_class_counts.tsv

```
reproducibility_class n_lineages
highly_robust 1
moderately_robust 7
method_sensitive 4
trim_sensitive 0
not_reproducible 6
```

#### S10.9 tier_asymmetry_tests.tsv

```
test_id scope outcome_variable core_value catalytic_value conditioning_set null_model test_statistic
    p_value_one_sided interpretation
primary_confirmed_allocation_hypergeometric primary_pathway_only confirmed_branches 11 3 total_confirmed
    =14; total_candidates=17; core_candidates=12; catalytic_candidates=5 confirmations are exchangeable
    among exploratory candidates irrespective of tier core_confirmed=11 (of 14) 0.191176 directional
    enrichment in core but weak-to-moderate evidence under small sample
primary_gene_label_permutation_confirmed primary_pathway_only confirmed_branches 11 3 4 genes total;
    exactly 2 genes assigned to core in each permutation (6 equally likely assignments) tier labels are
    exchangeable across the four genes observed_core_sum=11; max_permutation_sum=11 0.166667 observed
    split is extreme among six assignments but exact p remains non-small because of limited gene count
six_pathway_directional_sign_test six_pathway_sensitivity pathway_level_direction(core>catalytic) 4 1 non-
    tie pathways only (ties excluded by design) core and catalytic are equally likely to exceed each
    other per pathway (p=0.5) core_wins=4 of 5 non-tie pathways 0.187500 directional tendency favors core
     in sensitivity analysis, but statistical support is limited
```

#### S10.10 lineage_recurrence.tsv

```
canonical_lineage lineage n_genes_confirmed genes CLIPA8 CLIPB14 CLIPB15 SPCLIP1
cruzii A. cruzii 2 CLIPA8,SPCLIP1 1 0 0 1
funestus A. funestus 2 CLIPA8,SPCLIP1 1 0 0 1
maculipalpis A. maculipalpis 2 CLIPA8,CLIPB14 1 1 0 0
minimus A. minimus 2 CLIPB14,SPCLIP1 0 1 0 1
moucheti A. moucheti 2 CLIPA8,CLIPB15 1 0 1 0
christyi A. christyi 1 CLIPA8 1 0 0 0
```

```
culicifacies A. culicifacies 1 SPCLIP1 0 0 0 1
darlingi A. darlingi 1 CLIPA8 1 0 0 0
melas A. melas 1 SPCLIP1 0 0 0 1
```

#### S10.11 meme_sites_primary.tsv

```
gene query_id site meme_p overlaps_any_confirmed_beb_pp95
CLIPA8 AGAP010731 8 0.04586714233804612 False
CLIPA8 AGAP010731 10 0.01488751104012664 False
CLIPA8 AGAP010731 11 0.0400034821111771 True
CLIPA8 AGAP010731 12 0.004907487240129904 False
CLIPA8 AGAP010731 19 0.007439373452649001 False
CLIPA8 AGAP010731 21 0.02800778421484917 False
CLIPA8 AGAP010731 39 2.198806963171762e-07 False
CLIPA8 AGAP010731 67 0.03612233754458238 True
CLIPA8 AGAP010731 76 0.002017955147312622 False
CLIPA8 AGAP010731 192 0.03302887226694751 False
CLIPB14 AGAP010833 7 0.04532495998493336 False
CLIPB14 AGAP010833 9 0.02603406941329334 False
CLIPB14 AGAP010833 57 0.03941533579419987 False
CLIPB14 AGAP010833 74 0.01922020420003878 False
CLIPB14 AGAP010833 83 0.02822043230953564 False
CLIPB14 AGAP010833 244 0.01826357048042737 False
CLIPB14 AGAP010833 280 0.03031792871274142 False
CLIPB14 AGAP010833 299 0.01658420022285367 False
CLIPB15 AGAP009844 33 0.04948104513405882 False
CLIPB15 AGAP009844 56 0.0189466236917667 False
CLIPB15 AGAP009844 146 0.001917611276166187 True
CLIPB15 AGAP009844 151 0.008072513190641506 False
CLIPB15 AGAP009844 186 0.01807027020187035 False
CLIPB15 AGAP009844 216 0.003609241631772786 True
SPCLIP1 AGAP028725 13 0.02555433618806302 False
SPCLIP1 AGAP028725 15 0.007787321059072538 False
SPCLIP1 AGAP028725 16 0.003063517530748627 False
SPCLIP1 AGAP028725 20 0.001283870542288956 False
SPCLIP1 AGAP028725 24 0.0408923456357948 False
SPCLIP1 AGAP028725 63 0.01164007203188111 False
SPCLIP1 AGAP028725 76 0.04870788716679242 False
SPCLIP1 AGAP028725 77 0.01137862992208116 False
SPCLIP1 AGAP028725 108 0.0220350377511217 False
SPCLIP1 AGAP028725 139 0.001102796810259599 False
SPCLIP1 AGAP028725 147 0.009657774272807118 False
SPCLIP1 AGAP028725 336 0.002181299385119684 True
```

#### S10.12 pathway_robustness_summary.tsv

```
pathway method trim_state status status_by_gene total_confirmed_branches core_confirmed_branches
    catalytic_confirmed_branches total_absrel_candidates core_absrel_candidates
    catalytic_absrel_candidates branch_level_interpretable_all_genes site_level_interpretable_all_genes
    interpretation_note
babappalign_raw babappalign raw partial_meme_failure CLIPA8:hyphy_failed;CLIPB14:ok;CLIPB15:ok;SPCLIP1:ok
    11 8 3 16 10 6 True False Branch-level metrics interpretable across genes; MEME/site-level metrics
    partially missing. Directional outcome: core > catalytic. Non-ok status in: CLIPA8.
babappalign_clipkit babappalign clipkit ok CLIPA8:ok;CLIPB14:ok;CLIPB15:ok;SPCLIP1:ok 14 11 3 17 12 5 True
     True Fully comparable across genes. Directional outcome: core > catalytic.
mafft_raw mafft raw partial_meme_failure CLIPA8:hyphy_failed;CLIPB14:ok;CLIPB15:ok;SPCLIP1:ok 8 5 3 9 6 3
    True False Branch-level metrics interpretable across genes; MEME/site-level metrics partially missing
    . Directional outcome: core > catalytic. Non-ok status in: CLIPA8.
mafft_clipkit mafft clipkit ok CLIPA8:ok;CLIPB14:ok;CLIPB15:ok;SPCLIP1:ok 4 2 2 5 3 2 True True Fully
    comparable across genes. Directional outcome: core = catalytic.
prank_raw prank raw ok CLIPA8:ok;CLIPB14:ok;CLIPB15:ok;SPCLIP1:ok 5 3 2 5 3 2 True True Fully comparable
    across genes. Directional outcome: core > catalytic.
```

```
prank_clipkit prank clipkit partial_meme_failure CLIPA8:hyphy_failed;CLIPB14:ok;CLIPB15:ok;SPCLIP1:ok 3 1
    2 3 1 2 True False Branch-level metrics interpretable across genes; MEME/site-level metrics partially
     missing. Directional outcome: core < catalytic. Non-ok status in: CLIPA8.
```

#### S10.13 primary_normalized_signal_summary.tsv

```
gene tier final_codon_sites absrel_candidates codeml_confirmed beb_sites_branch_summed_pp95
    confirmations_per_tested_branch confirmed_branches_per_100_codons beb_codons_per_100_codons
CLIPA8 core 331 6 6 23 1.0000 1.8127 6.9486
CLIPB14 catalytic 308 3 2 3 0.6667 0.6494 0.9740
CLIPB15 catalytic 307 2 1 2 0.5000 0.3257 0.6515
SPCLIP1 core 347 6 5 20 0.8333 1.4409 5.7637
```

#### S10.14 reproducibility_lineage_table.tsv

```
gene query_id tier canonical_lineage display_lineage replication_count reproducibility_class
    pathways_present methods_present trim_states_present
CLIPB15 AGAP009844 catalytic moucheti A. moucheti 6 highly_robust babappalign_clipkit,babappalign_raw,
    mafft_clipkit,mafft_raw,prank_clipkit,prank_raw babappalign,mafft,prank clipkit,raw
CLIPA8 AGAP010731 core funestus A. funestus 4 moderately_robust babappalign_clipkit,babappalign_raw,
    mafft_raw,prank_raw babappalign,mafft,prank clipkit,raw
CLIPA8 AGAP010731 core maculipalpis A. maculipalpis 4 moderately_robust babappalign_clipkit,
    babappalign_raw,mafft_clipkit,mafft_raw babappalign,mafft clipkit,raw
CLIPA8 AGAP010731 core cruzii A. cruzii 3 moderately_robust babappalign_clipkit,babappalign_raw,mafft_raw
    babappalign,mafft clipkit,raw
CLIPA8 AGAP010731 core moucheti A. moucheti 3 moderately_robust babappalign_clipkit,prank_clipkit,
    prank_raw babappalign,prank clipkit,raw
CLIPB14 AGAP010833 catalytic minimus A. minimus 4 moderately_robust babappalign_clipkit,babappalign_raw,
    mafft_raw,prank_clipkit babappalign,mafft,prank clipkit,raw
SPCLIP1 AGAP028725 core minimus A. minimus 4 moderately_robust babappalign_clipkit,babappalign_raw,
    mafft_clipkit,mafft_raw babappalign,mafft clipkit,raw
SPCLIP1 AGAP028725 core culicifacies A. culicifacies 3 moderately_robust babappalign_clipkit,
    babappalign_raw,prank_raw babappalign,prank clipkit,raw
CLIPA8 AGAP010731 core christyi A. christyi 2 method_sensitive babappalign_clipkit,babappalign_raw
    babappalign clipkit,raw
CLIPB15 AGAP009844 catalytic aquasalis A. aquasalis 2 method_sensitive mafft_clipkit,mafft_raw mafft
    clipkit,raw
SPCLIP1 AGAP028725 core cruzii A. cruzii 2 method_sensitive babappalign_clipkit,babappalign_raw
    babappalign clipkit,raw
SPCLIP1 AGAP028725 core funestus A. funestus 2 method_sensitive babappalign_clipkit,babappalign_raw
    babappalign clipkit,raw
CLIPA8 AGAP010731 core darlingi A. darlingi 1 not_reproducible babappalign_clipkit babappalign clipkit
CLIPB14 AGAP010833 catalytic atroparvus A. atroparvus 1 not_reproducible babappalign_raw babappalign raw
CLIPB14 AGAP010833 catalytic gambiae A. gambiae 1 not_reproducible prank_raw prank raw
CLIPB14 AGAP010833 catalytic maculipalpis A. maculipalpis 1 not_reproducible babappalign_clipkit
    babappalign clipkit
SPCLIP1 AGAP028725 core aquasalis A. aquasalis 1 not_reproducible mafft_raw mafft raw
SPCLIP1 AGAP028725 core melas A. melas 1 not_reproducible babappalign_clipkit babappalign clipkit
```

#### S10.15 canonicalization_audit.tsv

```
gene query_id tier pathway method trim_state status raw_foreground_branch canonical_lineage
    display_lineage is_query_label is_confirmed_bh_0.05
CLIPA8 AGAP010731 core babappalign_raw babappalign raw hyphy_failed
    AFUN2_002201_R2179_Anopheles_funestus_AfunGA1 funestus A. funestus False True
CLIPA8 AGAP010731 core babappalign_raw babappalign raw hyphy_failed
    ACRUBR1_007112_R7575_Anopheles_cruzii_AcruBR1 cruzii A. cruzii False True
CLIPA8 AGAP010731 core babappalign_raw babappalign raw hyphy_failed
    AMOUCM1_004393_R4353_Anopheles_moucheti_AmouCM1 moucheti A. moucheti False False
```

```
CLIPA8 AGAP010731 core babappalign_raw babappalign raw hyphy_failed
    ACHR004918_RA_Anopheles_christyi_ACHKN1017 christyi A. christyi False True
CLIPA8 AGAP010731 core babappalign_raw babappalign raw hyphy_failed
    AMAC_004066_R3862_Anopheles_maculipalpis_AmacGA1 maculipalpis A. maculipalpis False True
CLIPA8 AGAP010731 core babappalign_clipkit babappalign clipkit ok
    AFUN2_002201_R2179_Anopheles_funestus_AfunGA1 funestus A. funestus False True
CLIPA8 AGAP010731 core babappalign_clipkit babappalign clipkit ok
    AMOUCM1_004393_R4353_Anopheles_moucheti_AmouCM1 moucheti A. moucheti False True
CLIPA8 AGAP010731 core babappalign_clipkit babappalign clipkit ok
    ACRUBR1_007112_R7575_Anopheles_cruzii_AcruBR1 cruzii A. cruzii False True
CLIPA8 AGAP010731 core babappalign_clipkit babappalign clipkit ok
    AMAC_004066_R3862_Anopheles_maculipalpis_AmacGA1 maculipalpis A. maculipalpis False True
CLIPA8 AGAP010731 core babappalign_clipkit babappalign clipkit ok
    ACHR004918_RA_Anopheles_christyi_ACHKN1017 christyi A. christyi False True
CLIPA8 AGAP010731 core babappalign_clipkit babappalign clipkit ok
    ADAR2_010765_R17226_Anopheles_darlingi_AdarGF1 darlingi A. darlingi False True
CLIPA8 AGAP010731 core mafft_raw mafft raw hyphy_failed AFUN021726_RA_Anopheles_funestus_FUMOZ funestus A.
     funestus False True
CLIPA8 AGAP010731 core mafft_raw mafft raw hyphy_failed AMAC_004066_R3862_Anopheles_maculipalpis_AmacGA1
    maculipalpis A. maculipalpis False True
CLIPA8 AGAP010731 core mafft_raw mafft raw hyphy_failed ACRUBR1_007112_R7575_Anopheles_cruzii_AcruBR1
    cruzii A. cruzii False True
CLIPA8 AGAP010731 core mafft_raw mafft raw hyphy_failed AMOUCM1_004393_R4353_Anopheles_moucheti_AmouCM1
    moucheti A. moucheti False False
CLIPA8 AGAP010731 core mafft_clipkit mafft clipkit ok AMAC_004066_R3862_Anopheles_maculipalpis_AmacGA1
    maculipalpis A. maculipalpis False True
CLIPA8 AGAP010731 core prank_raw prank raw ok AFUN021726_RA_Anopheles_funestus_FUMOZ funestus A. funestus
    False True
CLIPA8 AGAP010731 core prank_raw prank raw ok AMOUCM1_004393_R4353_Anopheles_moucheti_AmouCM1 moucheti A.
    moucheti False True
CLIPA8 AGAP010731 core prank_clipkit prank clipkit hyphy_failed
    AMOUCM1_004393_R4353_Anopheles_moucheti_AmouCM1 moucheti A. moucheti False True
CLIPB14 AGAP010833 catalytic babappalign_raw babappalign raw ok
    AMAC_000985_R425_Anopheles_maculipalpis_AmacGA1 maculipalpis A. maculipalpis False False
CLIPB14 AGAP010833 catalytic babappalign_raw babappalign raw ok
    ADAR2_004445_R6063_Anopheles_darlingi_AdarGF1 darlingi A. darlingi False False
CLIPB14 AGAP010833 catalytic babappalign_raw babappalign raw ok AMIN006979_RA_Anopheles_minimus_MINIMUS1
    minimus A. minimus False True
CLIPB14 AGAP010833 catalytic babappalign_raw babappalign raw ok AATE006733_RA_Anopheles_atroparvus_EBRO
    atroparvus A. atroparvus False True
CLIPB14 AGAP010833 catalytic babappalign_clipkit babappalign clipkit ok
    AMAC_000985_R425_Anopheles_maculipalpis_AmacGA1 maculipalpis A. maculipalpis False True
CLIPB14 AGAP010833 catalytic babappalign_clipkit babappalign clipkit ok
    ADAR2_004445_R6063_Anopheles_darlingi_AdarGF1 darlingi A. darlingi False False
CLIPB14 AGAP010833 catalytic babappalign_clipkit babappalign clipkit ok
    AMIN006979_RA_Anopheles_minimus_MINIMUS1 minimus A. minimus False True
CLIPB14 AGAP010833 catalytic mafft_raw mafft raw ok AMIN006979_RA_Anopheles_minimus_MINIMUS1 minimus A.
    minimus False True
CLIPB14 AGAP010833 catalytic prank_raw prank raw ok AGAP010833 gambiae A. gambiae True True
CLIPB14 AGAP010833 catalytic prank_clipkit prank clipkit ok AMIN006979_RA_Anopheles_minimus_MINIMUS1
    minimus A. minimus False True
CLIPB15 AGAP009844 catalytic babappalign_raw babappalign raw ok
    AMOUCM1_000648_R17196_Anopheles_moucheti_AmouCM1 moucheti A. moucheti False True
CLIPB15 AGAP009844 catalytic babappalign_raw babappalign raw ok AMIN009569_RA_Anopheles_minimus_MINIMUS1
    minimus A. minimus False False
CLIPB15 AGAP009844 catalytic babappalign_clipkit babappalign clipkit ok
    AMOUCM1_000648_R17196_Anopheles_moucheti_AmouCM1 moucheti A. moucheti False True
CLIPB15 AGAP009844 catalytic babappalign_clipkit babappalign clipkit ok
    AMIN009569_RA_Anopheles_minimus_MINIMUS1 minimus A. minimus False False
CLIPB15 AGAP009844 catalytic mafft_raw mafft raw ok AMOUCM1_000648_R17196_Anopheles_moucheti_AmouCM1
    moucheti A. moucheti False True
CLIPB15 AGAP009844 catalytic mafft_raw mafft raw ok AAQUA_001018_R21657_Anopheles_aquasalis aquasalis A.
    aquasalis False True
CLIPB15 AGAP009844 catalytic mafft_clipkit mafft clipkit ok
    AMOUCM1_000648_R17196_Anopheles_moucheti_AmouCM1 moucheti A. moucheti False True
```

```
CLIPB15 AGAP009844 catalytic mafft_clipkit mafft clipkit ok AAQUA_001018_R21657_Anopheles_aquasalis
     aquasalis A. aquasalis False True
CLIPB15 AGAP009844 catalytic prank_raw prank raw ok AMOUCM1_000648_R17196_Anopheles_moucheti_AmouCM1
     moucheti A. moucheti False True
CLIPB15 AGAP009844 catalytic prank_clipkit prank clipkit ok
     AMOUCM1_000648_R17196_Anopheles_moucheti_AmouCM1 moucheti A. moucheti False True
SPCLIP1 AGAP028725 core babappalign_raw babappalign raw ok ACUA011631_RA_Anopheles_culicifacies
     culicifacies A. culicifacies False True
SPCLIP1 AGAP028725 core babappalign_raw babappalign raw ok AMEC000258_RA_Anopheles_melas_CM1001059_A melas
      A. melas False False
SPCLIP1 AGAP028725 core babappalign_raw babappalign raw ok AFUN2_012038_R21717_Anopheles_funestus_AfunGA1
     funestus A. funestus False True
SPCLIP1 AGAP028725 core babappalign_raw babappalign raw ok ACRUBR1_005064_R5104_Anopheles_cruzii_AcruBR1
     cruzii A. cruzii False True
SPCLIP1 AGAP028725 core babappalign_raw babappalign raw ok AMIN004295_RA_Anopheles_minimus_MINIMUS1
     minimus A. minimus False True
SPCLIP1 AGAP028725 core babappalign_clipkit babappalign clipkit ok ACUA011631_RA_Anopheles_culicifacies
     culicifacies A. culicifacies False True
SPCLIP1 AGAP028725 core babappalign_clipkit babappalign clipkit ok
     AFUN2_012038_R21717_Anopheles_funestus_AfunGA1 funestus A. funestus False True
SPCLIP1 AGAP028725 core babappalign_clipkit babappalign clipkit ok
     AMEC000258_RA_Anopheles_melas_CM1001059_A melas A. melas False True
SPCLIP1 AGAP028725 core babappalign_clipkit babappalign clipkit ok
     ACRUBR1_005064_R5104_Anopheles_cruzii_AcruBR1 cruzii A. cruzii False True
SPCLIP1 AGAP028725 core babappalign_clipkit babappalign clipkit ok
     AMIN004295_RA_Anopheles_minimus_MINIMUS1 minimus A. minimus False True
SPCLIP1 AGAP028725 core babappalign_clipkit babappalign clipkit ok
     AMAM010672_RA_Anopheles_maculatus_maculatus3 maculatus A. maculatus False False
SPCLIP1 AGAP028725 core mafft_raw mafft raw ok AMIN004295_RA_Anopheles_minimus_MINIMUS1 minimus A. minimus
      False True
SPCLIP1 AGAP028725 core mafft_raw mafft raw ok AAQUA_009556_R14763_Anopheles_aquasalis aquasalis A.
     aquasalis False True
SPCLIP1 AGAP028725 core mafft_clipkit mafft clipkit ok AMIN004295_RA_Anopheles_minimus_MINIMUS1 minimus A.
      minimus False True
SPCLIP1 AGAP028725 core mafft_clipkit mafft clipkit ok AMEC000258_RA_Anopheles_melas_CM1001059_A melas A.
     melas False False
SPCLIP1 AGAP028725 core prank_raw prank raw ok ACUA011631_RA_Anopheles_culicifacies culicifacies A.
     culicifacies False True
```

#### S11. Interpretive rule used in revision

The revised manuscript prioritizes robustness-aware interpretation:

- recurring patterns are reported as empirical support;
- pathway-sensitive outcomes are explicitly retained;
- non-significant asymmetry tests are acknowledged directly;
- no decisive pathway-level confirmation is claimed.

### Supplementary Figures

#### S12. Figure list used in manuscript package

- `fig_babappasnake_workflow_architecture.pdf`
- `fig_pathway_tiered_model.pdf`

- `fig_primary_branch_site_summary.pdf`
- `fig_lineage_recurrence_heatmap.pdf`
- `fig_alignment_robustness_heatmap.pdf`
- `fig_pathway_tier_summary.pdf`
- `fig_branch_reproducibility_matrix.pdf`

#### S13. Embedded workflow and case-study figures (full panels)

All key workflow and mosquito-case-study figures are embedded below so this supplementary document is self-sufficient for interpretation without external figure lookup.

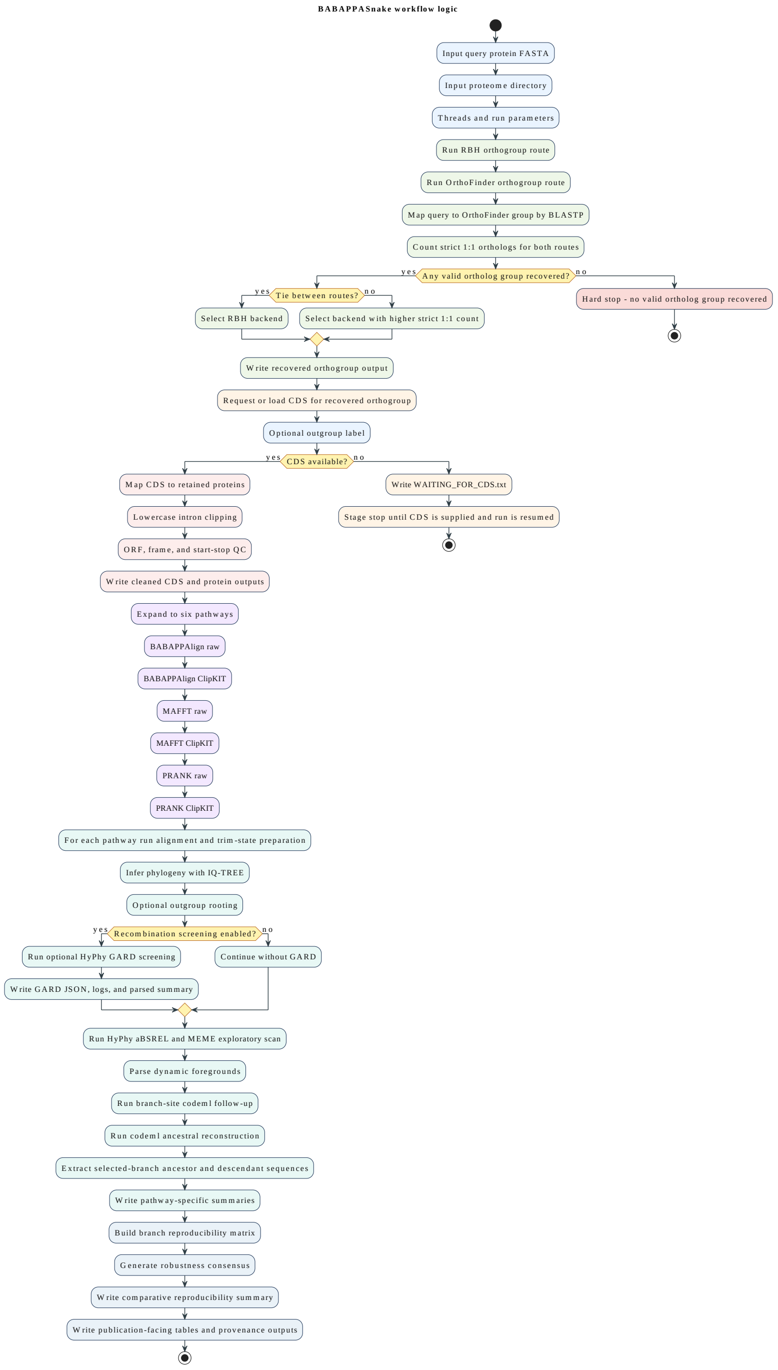


Figure 1: Decision-aware BABAPPASnake architecture with backend selection, CDS stage gating, method×trim pathway branching, repeated per-pathway inference, and final cross-pathway synthesis/provenance outputs.

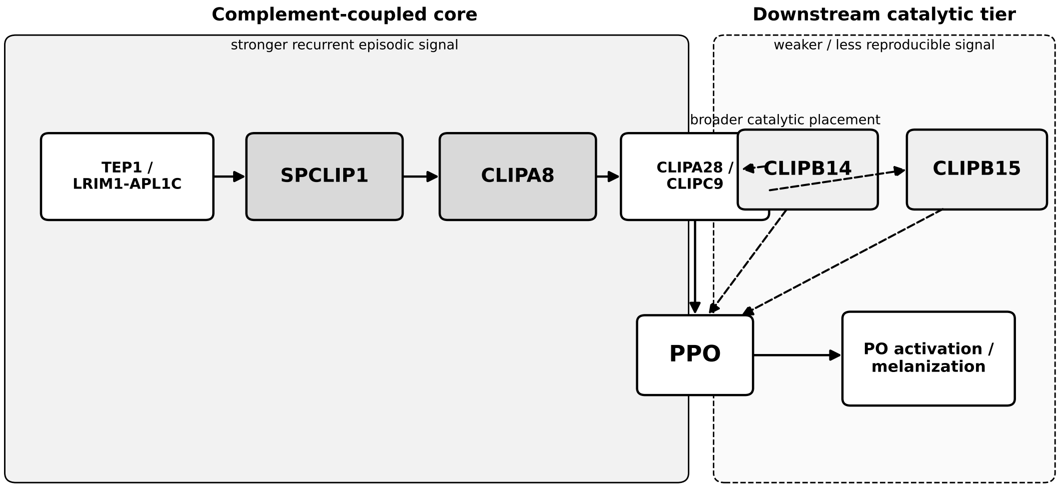


Figure 2: Mosquito melanization module layout used in the empirical demonstration, showing complement-coupled core tier and catalytic tier organization.

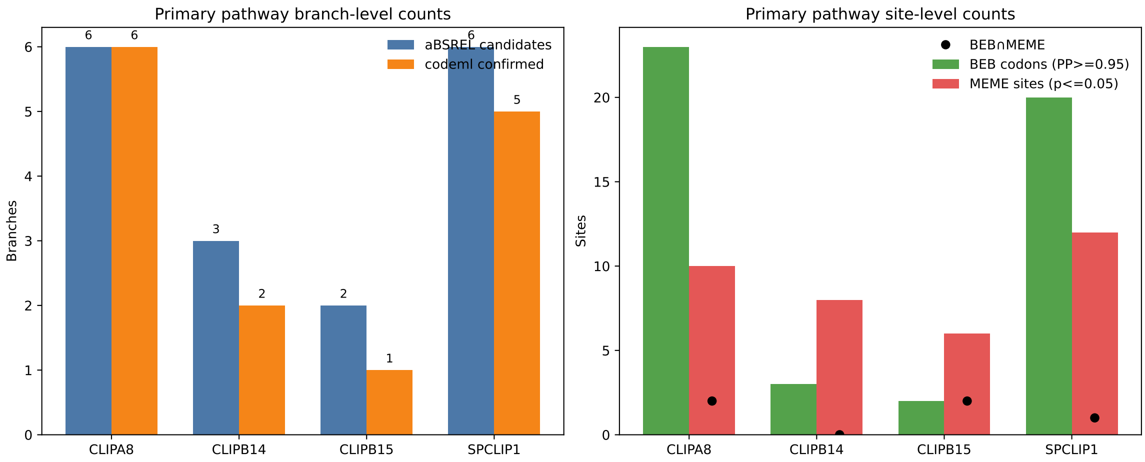


Figure 3: Primary pathway branch-site summary for the four-gene mosquito demonstration, including exploratory and follow-up-positive lineage patterns.

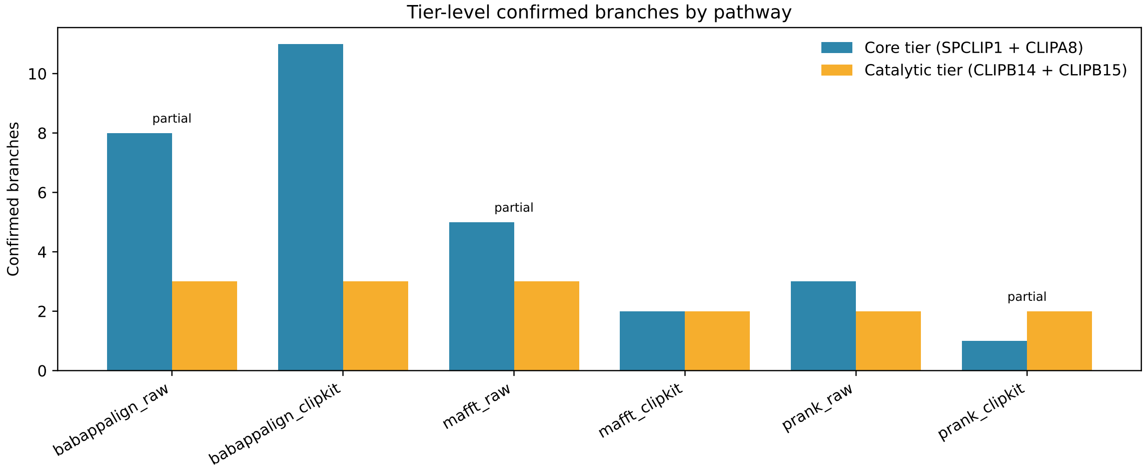


Figure 4: Bar-plot style pathway-tier summary across method×trim pathways, showing core-vs-catalytic follow-up-positive branch counts.

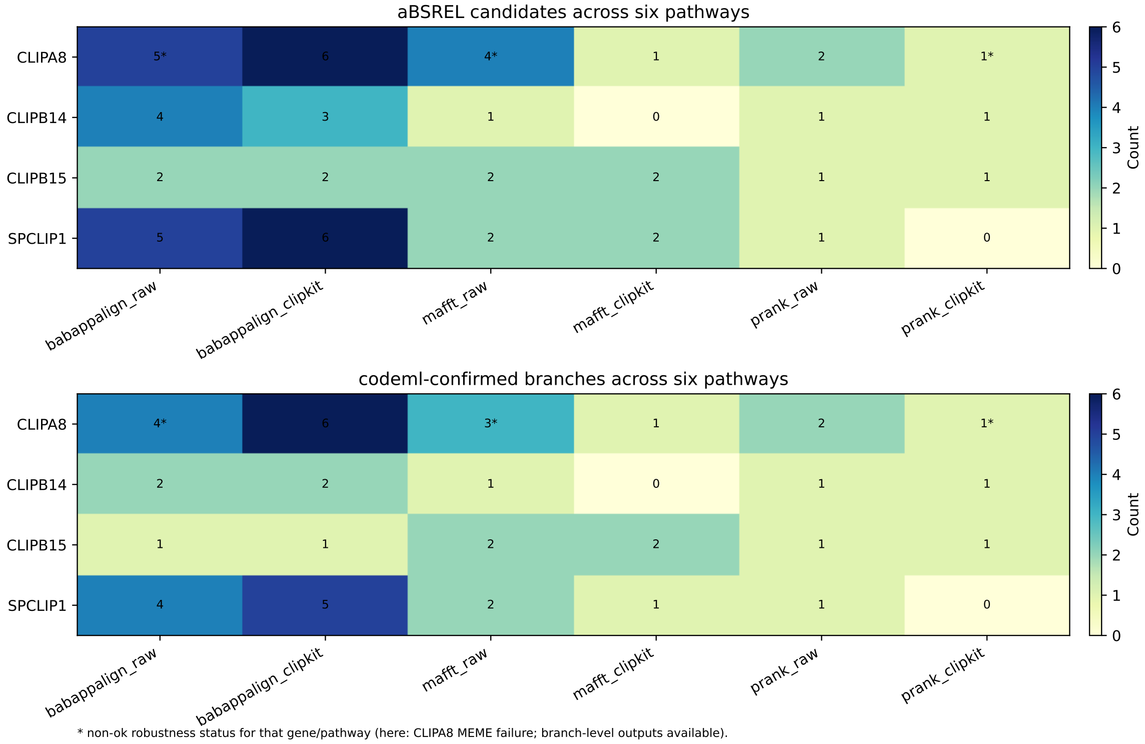


Figure 5: Alignment/trim robustness heatmap across six pathways, highlighting reproducibility and pathway sensitivity patterns.

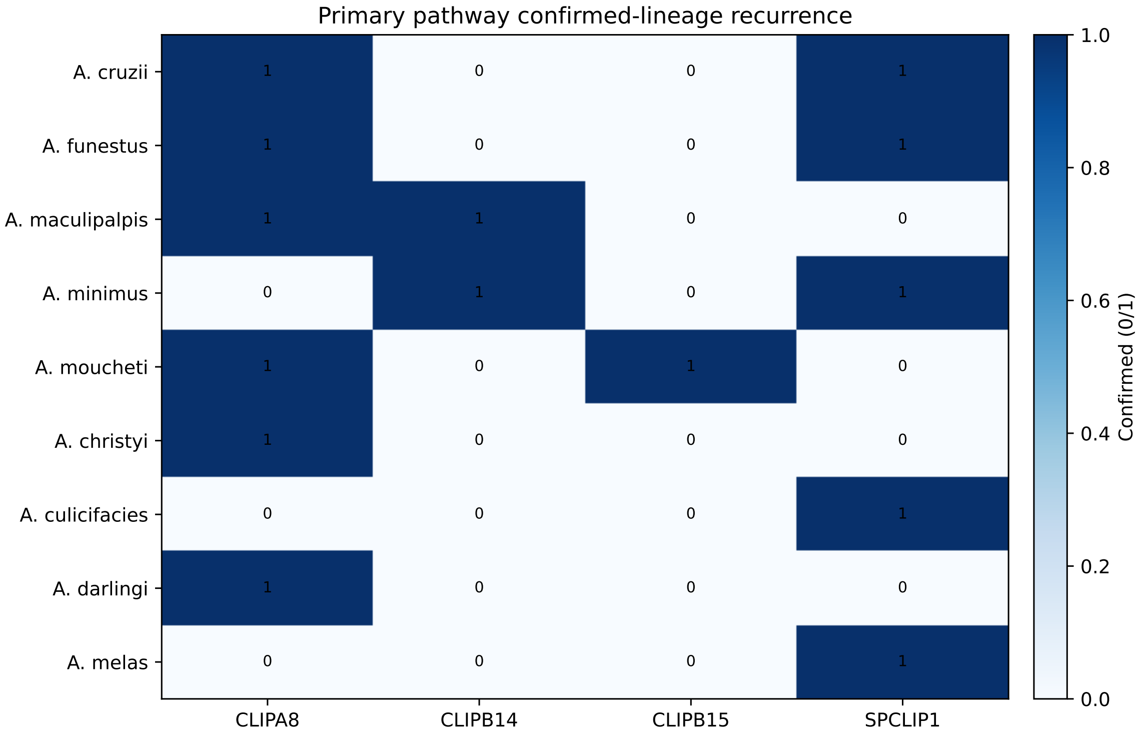


Figure 6: Lineage recurrence heatmap summarizing repeated versus pathway-restricted branch signals in the mosquito case study.

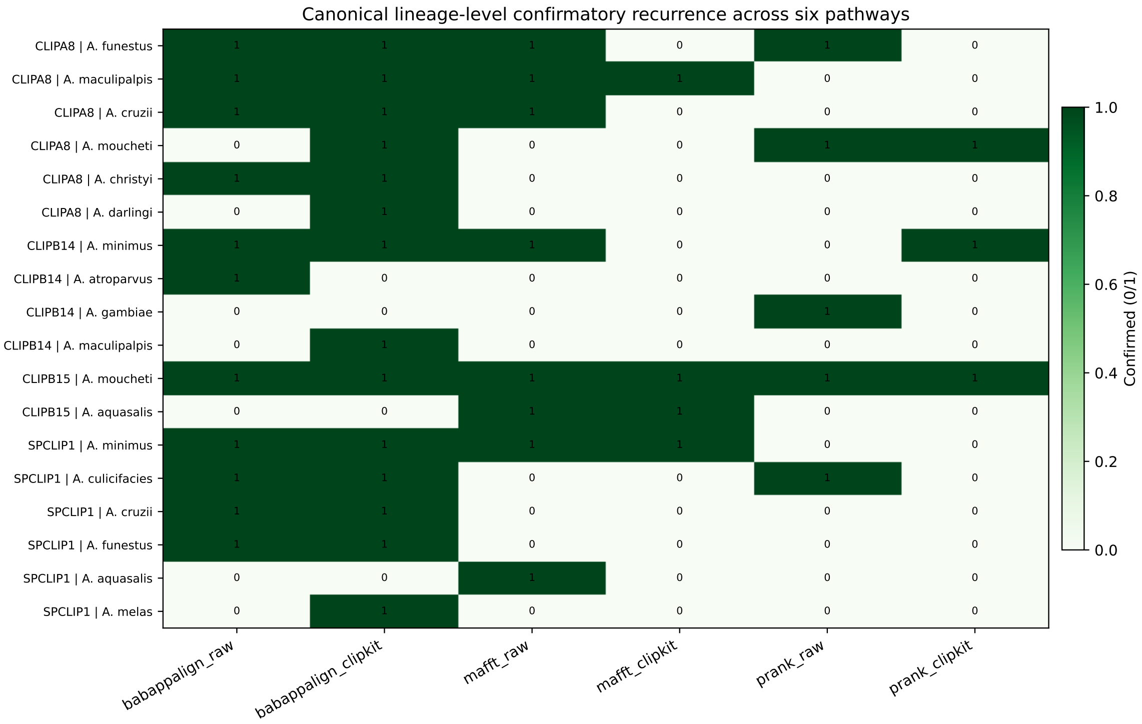


Figure 7: Branch reproducibility matrix across method×trim pathways used for robustness class assignment and comparative interpretation.

### File and Archive Guide

#### S14. Where each reproducibility layer is located

- **Software/workflow logic**: BABAPPASnake repository.
- **Run-specific outputs**: archived case-study run directories and generated tables.
- **Manuscript companion**: this supplementary file, which reorganizes the same outputs for reader navigation.

#### S15. Limitation statement (supplementary)

No broad cross-pipeline benchmarking is introduced in this revision. The manuscript is intentionally positioned as a workflow introduction with empirical demonstration; broader benchmarking remains future work.
